# Controlling swelling and release of hyaluronic acid during aqueous storage by *in situ* cross-linking during spray drying with alginate

**DOI:** 10.1101/679589

**Authors:** Dana E. Wong, Julia C. Cunniffe, Herbert B. Scher, Tina Jeoh

## Abstract

The success of hyaluronic acid in over-the-counter cosmetics has been limited by its poor storage stability in aqueous environments due to premature swelling and hydrolysis. Here, hyaluronic acid was prepared in dry microparticles, encapsulated by spray-drying in patented *in situ* calcium cross-linked alginate microcapsules (CLAMs) to minimize swelling and release in aqueous formulations. CLAMs prepared with 61% (d.b.) hyaluronic acid (HA-CLAMs) demonstrated restricted plumping, limited water absorption capacity, and reduced leaching; retaining up to 49 % hyaluronic acid after 2 hrs in water. A new method using chelated soluble calcium resulted in particles with significantly improved hyaluronic acid retention in water. ‘Chelate HA-CLAMs’ exhibited nearly full retention of hyaluronic acid over 2 hr incubation in water, and remained visibly insoluble after 1 year of storage in water at 4^°^C. Successful hyaluronic acid retention in CLAMs is likely due to the ability of hyaluronic acid to participate in calcium cross-linking.

## 1. INTRODUCTION

Hyaluronic acid or hyaluronan, is an important naturally occurring anionic glycosaminoglycan biopolymer in the body. Hyaluronic acid is found in cell membranes, as synovial fluid lubricant between joints, as the main component in ocular fills, and is a major component of skin (Song et al., 2009). Hyaluronic acid is composed of regularly ordered β-D-glucuronic acid and N-acetyl-β-D-glucosamine linked by alternating β(1**→**4) and β(1**→**3) glycosidic bonds. The proximity of the amide and carboxylic acid groups within each repeating unit facilitates tight water associations with the polymer such that hyaluronic acid is a widely documented humectant (Caspersen et al., 2014, Beasley et al., 2009). The polyuronic acid molecular structure avails repeating carboxylic acid groups along the biopolymer that allows for extensive electrostatic interactions with ions. Hyaluronic acid matrices are prevalently used in a range of medical applications such as arthritic injections, wound healing, and drug delivery because of its abundance and natural biocompatibility (Travan et al., 2016, Vasi et al., 2014, Collins and Birkinshaw, 2013, Iskandar et al., 2009). Hyaluronic acid can improve the biocompatibility of mixed polymer matrices due to its natural presence in the human body (Jou et al., 2007), and can be used in tandem with other biopolymers like chitosan and alginate (Almalik et al., 2013, Nath et al., 2015, Gao et al., 2014). Previously, hyaluronic acid use in cosmetics was limited to surgical fillers and semi-permanent injections (Rohrich et al., 2007, Juhász and Marmur, 2015, Ho and Jagdeo, 2015); more recently, hyaluronic acid use has expanded into over-the-counter personal care and cosmetics products (Ammala, 2013, Beasley et al., 2009, Janiš et al., 2017). The rise of topical cosmetics applications for hyaluronic acid is due to its production abundance, efficacy, and biocompatibility.

Delivering active hyaluronic acid from aqueous formulations to dermal sites is challenging because of its extreme hydroscopic nature. The success of hyaluronic acid in over-the-counter cosmetics has been limited by its high cost and poor storage stability in aqueous environments due to premature swelling and hydrolysis. Premature swelling of hyaluronic acid during storage prior to application can compromise absorption if the hydrated polymer is larger than the size of a skin pore (Pilkington et al., 2015). Furthermore, pure hyaluronic acid in aqueous cosmetics are susceptible to premature hydrolysis, which leads to lowered moisture retention activity on the skin. For example, Simulescu et al. found that hyaluronic acid of varying sizes were susceptible to 90% weight average molecular weight decreases when stored at room temperature for 2 months without antimicrobials or protectants. Refrigeration limited but did not prevent hydrolysis when in solution (Simulescu et al., 2016). Mondek found that although the overall hyaluronic acid polydispersity remained unchanged, degradation was attributed to temperature (Mondek et al., 2015). Molecular weight changes were observed at both room (25°C) and refrigerated (4°C) temperatures. For successful topical hyaluronic acid application, effective strategies for extended storage in aqueous formulations and controlled release on skin are clearly needed.

Many have explored encapsulation as a method to protect and deliver hyaluronic acid for medical applications. In a hyaluronic acid preparation for deep dermal drug delivery, Berkó et al. found that chemically cross-linking hyaluronic acid helped to maintain its desirable hydration effect. Even when smaller particles were produced due to the nature of the cross-linking method, there were limited changes to the overall rheolgical properties of hyaluronic acid. These small cross-linked hyaluronic acid particles were able to improve membrane diffusion and skin penetration compared to linear hyaluronic acid (Berkó et al., 2013).

Only a few have attempted to spray dry hyaluronic acid, all with final applications in pharmaceuticals (Iskandar et al., 2009, Huh et al., 2010). Cross-linked Alginate Microcapsules (CLAMs) provide a storage method that is capable of incorporating cargo without chemical modification into its matrix and prevents release in water. CLAMs produced by a patented, industrially-scalable, one-step method developed by our research group is pH mediated, where a feed suspension of cargo, encapsulant (alginate), calcium salt, weak acid, and volatile base are spray dried such that gelling, curing and drying occur *in situ*, Figure 1, (Jeoh-Zicari et al., 2017). In contrast, traditional methods for forming CLAMs require multiple time consuming steps that are prohibitively costly to scale up (Strobel et al., 2019a). Like hyaluronic acid, alginate is also a hydroscopic, polyuronic acid biopolymer with high solution viscosity (Draget and Taylor, 2011). The resulting CLAMs are water insoluble, maintain barrier properties in storage, and exhibit unique release characteristics.

**Figure 1:**
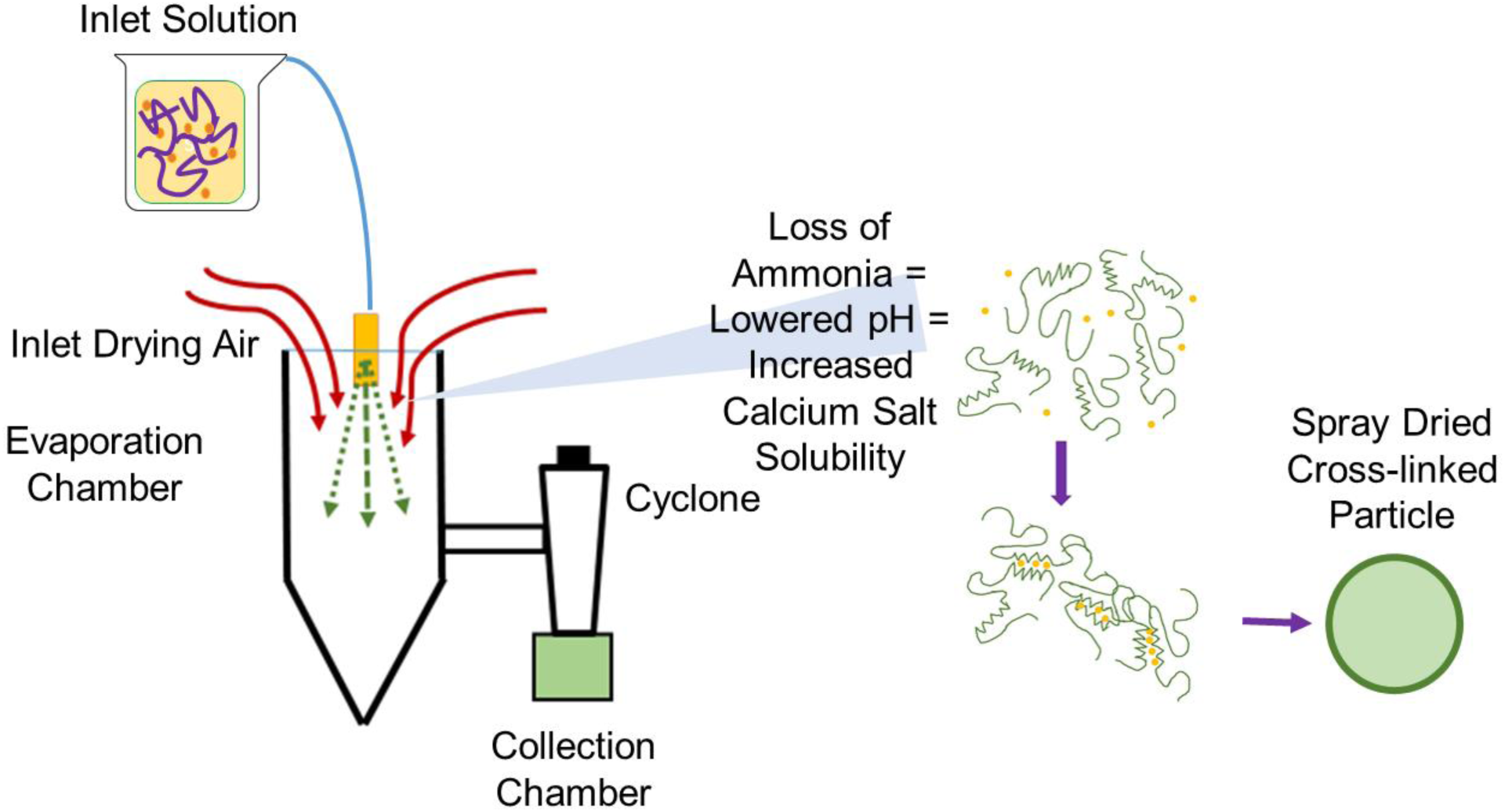
Schematic of patented technology for pH-mediated electrostatic cross-linking during spray-drying to produce dry, cross-linked microparticles.

In this study, the encapsulation of hyaluronic acid in CLAMs by spray drying was explored, with the goal of producing dry particles with limited swelling in aqueous storage. Hyaluronic acid was incorporated into the encapsulation matrix of alginate, and the resulting dry hyaluronic acid loaded CLAMs (HA-CLAM) were characterized. Additionally, an alternate CLAMs formulation utilizing pH responsive chelated calcium in the feed was explored. The results of these studies suggest new opportunities for cost-effective and industrially-scalable hyaluronic acid/CLAMs encapsulation matrices for drug delivery and tissue regeneration applications (Al-Sibani et al., 2017, Ekici et al., 2011, Ganesh et al., 2013).

## 2. EXPERIMENTAL

### 2.1 Materials

High viscosity (HV) sodium alginate (Hydagen 155P) and sodium hyaluronate/hyaluronan (36 kDa) was provided by BASF SE. Calcium phosphate, succinic acid, sodium citrate, glacial acetic acid, sodium carbonate, β-D-glucose, sodium hydroxide, hydrochloric acid, calcium carbonate, sodium citrate, phytic acid (inositol hexakisphosphate), sodium tetraborate, sulfuric acid, and ammonium hydroxide were purchased from Thermo Fisher. Schiff’s fuchsin sulfite reagent, sodium metabisulfate, periodic acid, calcium chloride, citric acid, low viscosity (LV) sodium alginate from brown algae (cat#A1112), and carbazole were purchased from Millipore Sigma. Ethanol (200 proof) was purchased from Koptek. Carbon tape and microscopy stands were purchased from Ted Pella. Ultrapure deionized water was sourced from a MilliQ 85/15 system (Millipore Sigma).

### 2.2 Methods

#### 2.2.1 Spray Dried Cross-linked Alginate Microcapsules (CLAMs)

CLAMs were formed with and without hyaluronic acid as cargo following previously published methods with some adjustments (Jeoh-Zicari et al., 2017, Strobel et al., 2017, Strobel et al., 2016, Strobel et al., 2019b). Spray dryer feed formulations using either low viscosity (LV) or high viscosity (HV) alginates at 0.5% (w/w) were adjusted to pH 5.4 and pH 7, respectively, by titrating succinic acid with ammonium hydroxide to maintain insoluble calcium hydrogen phosphate (as 0.0625% of the feed or 12.5% d.b. w/w of the final CLAMs). Succinic acid was included at one half the concentration of alginate, or 0.25% (w/w) of the inlet feed. Hyaluronic acid loaded CLAMs (HA-CLAMs) were prepared by introducing sodium hyaluronate to the feed solution during alginate hydration. HA-CLAMs formulations were prepared with 1.25 % (w/w) sodium hyaluronate resulting in 61% dry basis final hyaluronic acid content.

‘Chelate CLAMs’ were also formed with and without hyaluronic acid cargo. This variation on the methods outlined above used HV alginate at 0.5% (w/w) in the spray dryer feed formulation. The feed solution was prepared by titrating a solution of phytic acid (inositol hexakisphosphate) to pH 8.4 with ammonium hydroxide. Phytic acid was included in the feed formulation at 5 times the concentration of calcium chloride (0.625% of solution). Hyaluronic acid was incorporated to the chelate solution to achieve 61% dry basis final hyaluronic acid content to be directly compared to previous formulations.

Spray dried hyaluronic acid particles were formed with calcium hydrogen phosphate to assess the ability of hyaluronic acid to participate in ion-mediated cross-linking. Solutions were prepared with 2% (w/w) hyaluronic acid and twice the concentration of succinic acid. Succinic acid was titrated to pH 5.6 with ammonium hydroxide to prevent the solubility of calcium hydrogen phosphate which was included at 0.25% (w/w) in solution. All formulations were prepared at a 2:1 ratio of alginate or hyaluronic acid to succinic acid and an 8:1 ratio of polymer to calcium. All the variations of the CLAMs tested in this study are summarized in Table 1.

**Table 1:**
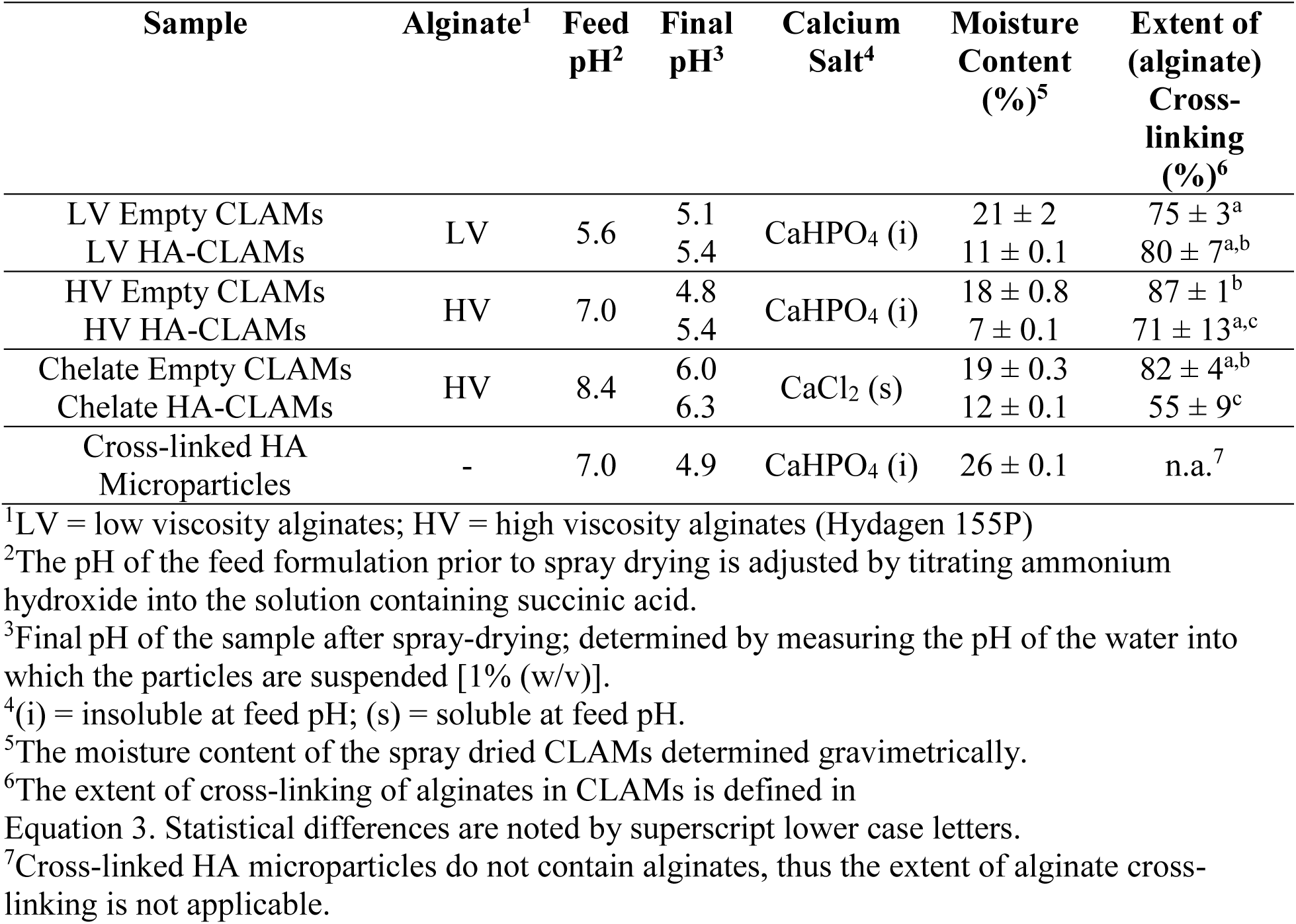
Formulations used in the microencapsulation of hyaluronic acid. Final product for all formulations were collected in the form of a dry, white powder.

| Sample | Alginate<sup>1</sup> | Feed pH<sup>2</sup> | Final pH<sup>3</sup> | Calcium Salt<sup>4</sup> | Moisture Content (%)<sup>5</sup> | Extent of (alginate) Cross-linking (%)<sup>6</sup> |
| --- | --- | --- | --- | --- | --- | --- |
| LV Empty CLAMs | LV | 5.6 | 5.1 | CaHPO<sub>4</sub> (i) | 21 ± 2 | 75 ± 3<sup>a</sup> |
| LV HA-CLAMs |  |  | 5.4 |  | 11 ± 0.1 | 80 ± 7<sup>a,b</sup> |
| HV Empty CLAMs | HV | 7.0 | 4.8 | CaHPO<sub>4</sub> (i) | 18 ± 0.8 | 87 ± 1<sup>b</sup> |
| HV HA-CLAMs |  |  | 5.4 |  | 7 ± 0.1 | 71 ± 13<sup>a,c</sup> |
| Chelate Empty CLAMs | HV | 8.4 | 6.0 | CaCl<sub>2</sub> (s) | 19 ± 0.3 | 82 ± 4<sup>a,b</sup> |
| Chelate HA-CLAMs |  |  | 6.3 |  | 12 ± 0.1 | 55 ± 9<sup>c</sup> |
| Cross-linked HA Microparticles | - | 7.0 | 4.9 | CaHPO<sub>4</sub> (i) | 26 ± 0.1 | n.a.<sup>7</sup> |

<sup>1</sup>LV = low viscosity alginates; HV = high viscosity alginates (Hydagen 155P)
<sup>2</sup>The pH of the feed formulation prior to spray drying is adjusted by titrating ammonium hydroxide into the solution containing succinic acid.
<sup>3</sup>Final pH of the sample after spray-drying; determined by measuring the pH of the water into which the particles are suspended [1% (w/v)].
<sup>4</sup>(i) = insoluble at feed pH; (s) = soluble at feed pH.
<sup>5</sup>The moisture content of the spray dried CLAMs determined gravimetrically.
<sup>6</sup>The extent of cross-linking of alginates in CLAMs is defined in Equation 3. Statistical differences are noted by superscript lower case letters.
<sup>7</sup>Cross-linked HA microparticles do not contain alginates, thus the extent of alginate cross-linking is not applicable.

CLAMs were produced in a benchtop spray dryer (B-290, Büchi, New Castle, DE) at an inlet temperature of 150°C, aspirator air flow of 35 m^3^/hr (maximum), feed pump at 20% of the maximum, and air nozzle flow at 40 mm. The resulting dried powders were stored in dessicators until analysis.

#### 2.2.2 Monitoring Hyaluronic Acid Release in Water

Hyaluronic acid release from HA-CLAMs in water was monitored over 120 min. Both empty CLAMs and HA-CLAMs were suspended at 1% (w/v) in water in separate tubes for each timed point. Samples were analyzed for hyaluronic acid release at 0, 15, 30, 45, 60, 90, and 120 min after continuous rotation at 25 rpm. Two minutes prior to the predetermined time, samples were centrifuged at 5000 rpm for 2 min to separate residual solid CLAMs from the supernatant. The supernatant was diluted 50 times in water and measured for soluble alginate and total uronic acid (alginates + hyaluronic acid) concentrations by the Periodic Schiff’s Fuchsin Assay and the Carbazole Assay, respectively. Hyaluronic acid released into solution at each time was calculated as:

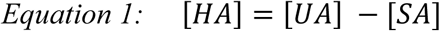

Where, [HA] is the concentration of hyaluronic acid (mg/mL), [UA] is the total concentration of uronic acids (mg/mL), and [SA] is the concentration of soluble alginate (mg/mL) released in the solution at the given time.

Hyaluronic acid released as a percentage of the total concentration of hyaluronic acid upon full release ([HA]full release) from the microcapsules at each time point was calculated as:

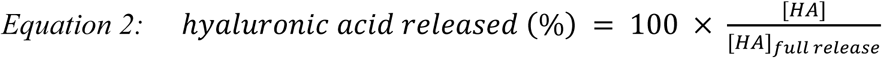

Empty CLAMs served as controls in all release measurements.

#### 2.2.3 Determining the extent of Cross-linking of CLAMs – Periodic Schiff’s Fuchsin Assay

CLAMs fully dissolve in chelating solutions that sequester ions and disrupt crosslinks but remain insoluble in non-chelating, aqueous solutions. Thus, the extent of alginate cross-linking in CLAMs is defined as the fraction of insoluble alginates when suspended in a non-chelating, aqueous solution (Strobel et al., 2017, Strobel et al., 2019b). To measure the extent of cross-linking, alginates solubilized from CLAMs suspended in DI water (non-chelating solution) or 100 mM sodium citrate buffer (chelating solution) are compared. CLAMs at 1% (w/w) were incubated for 2 h with end over end mixing in the respective solutions at room temperature, then centrifuged for 5 min at 200 rpm to separate the supernatant from any remaining insoluble CLAMs. The supernatant was diluted in water 50 times, from which 200 µL was mixed with 30 µL of a solution of periodic acid (4.67 % w/v) and acetic acid (0.67 % w/v) in microtiter plates and incubated at 37°C for 1 h. The Schiff’s Reagent was prepared as 66.67 % (v/v) solution of Schiff’s fuchsin sulfite in water with 100.2 mg sodium metabisulfate, and also incubated at 37°C for 1 h. After incubation of both the samples and the reagent, 30 µL of the Schiff’s Reagent was added to sample wells, the microtiter plate was wrapped in foil, and held at room temperature for 45 min to develop color. Sample absorbances were measured at 550 nm in a plate reader (BioTek Synergy 4) and compared to a standard curve of alginates ranging between 0 – 0.25 mg/mL to determine the concentration of alginates in each solution. Separate standard curves were generated for the low and high viscosity alginates. The extent of cross-linking of the CLAMs was calculated as follows:

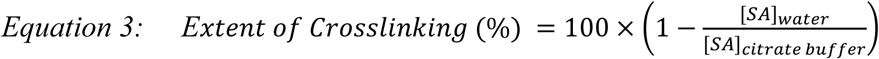

Where [SA]water is the concentration of solubilized alginates in DI water (mg/mL) and [SA]citrate buffer is the concentration of solubilized alginates in the sodium citrate buffer (mg/mL).

The Schiff’s reagent does not react with hyaluronic acid and is therefore specific to alginate quantitation when both alginates and hyaluronic acids are present in solution.

#### 2.2.4 Carbazole Assay for Uronic Acid Quantitation

The carbazole assay measures total uronic acids in solution (Bitter and Muir, 1962, Blumenkrantz and Asboe-Hansen, 1973, Cesaretti et al., 2003). The 50 times diluted samples (50 µL) of supernatant from the hyaluronic acid release experiments were combined with 200 µL of 25 mM sodium tetraborate in 72% sulfuric acid in a polymerase chain reaction (PCR) plate and heated at 100°C for 10 min in a thermocycler (Bio-Rad MyCycler 1709703). After cooling to room temperature, 50 µL of 0.125 mM carbazole in 200 proof ethanol was added to each well and incubated for an additional 10 min at 100°C. Sample absorbances were read at 550 nm in a plate reader (Biotek Synergy 4) when cooled to room temperature and compared to a standard curve of equal parts alginate and sodium hyaluronate a total concentrations of 0 – 100 µg/mL in water.

#### 2.2.5 Scanning Electron Microscopy (SEM)

Samples were adhered to stands with carbon tape and coated with 15 nm gold using a Cressington 108 Auto Coating System (Watford, UK). All micrographs of CLAMs were produced by a FEI/Philips XL 30 SFEG SEM or a FE-SEM Hitachi S-4100T. Unless indicated, micrographs were produced with an operating voltage of 5 kV. Representative micrographs from numerous captures at varying magnifications are presented. Micrographs were captured at the NEAT ORU Keck Spectral Imaging Facility and Advanced Materials Characterization and Testing Lab at the University of California, Davis.

#### 2.2.6 Quantifying the Water Absorption Capacity (WAC) and Plumping Ratio of the cross-linked microparticles in water

Water Absorption Capacity (WAC), the mass ratio of a hydrated product compared to its dry counterpart, indicates the extent to which the cross-linked microparticles (with and without hyaluronic acid) uptake water. The plumping ratio, i.e. the volume ratio of hydrated product compared to its dry counterpart, is a measure of the product volume change due to absorption of water. Plumping indicates the extent to which the cross-linked microparticles (with and without hyaluronic acid) swell upon water absorption.

Dry powder samples (Table 1) (0.1 g) were added to 2 g of DI water in 13 × 100 flat bottom glass test tubes and allowed to sit for 45 min at room temperature. After 45 min, the test tubes were centrifuged at 190 *g* for 3 min in a swinging bucket centrifuge. The separated supernatant was removed and weighed. The mass of the hydrated powder was determined from the difference between the mass of the dry powder and added water (2.1 g) and the mass of the supernatant. The hydrated product height was measured using calipers.

The WAC was calculated as follows:

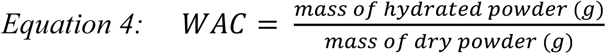

The height difference between dry and hydrated product was used to determine the plumping ratio in water.

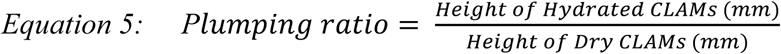

#### 2.2.7 Particle Sizing

Average particle diameter was measured using a Mastersizer 3000 (Malvern, UK) with isopropanol diluent to prevent swelling. Particles were submerged in isopropanol and sonicated for 10 min before measurement. The refractive index of the samples were compared to that of sugar (0.1). Data was collected as an average of 10 measurements.

#### 2.2.8 Statistics

Statistical analyses were conducted using Graphpad Prism software (v. 7.03, GraphPad Software, La Jolla, CA). Determination of statistical differences between sample means with a 95% CI (p<0.05) was done by One-Way ANOVA with Tukey’s pairwise comparisons when noted. All error values are reported as standard deviation calculated from a specified n equal to or greater than 3 when noted.

## 3. RESULTS AND DISCUSSION

### 3.1 Microencapsulation of hyaluronic acid in dry, cross-linked microparticles

The strategy of microencapsulating hyaluronic acid in cross-linked alginates was explored to minimize release and thus degradation of the polymer during storage in aqueous formulations. Microcapsules in this study were formed by a patented, one-step, industrially-scalable spray-drying method of in situ ion-mediated cross-linking of acidic polymers to form dry, cross-linked alginate microcapsules (CLAMs) (Jeoh-Zicari et al., 2012). The CLAMs formulation was varied (Table 1) to examine the role of alginates and the process to control the availability of calcium ions on the physical properties and hyaluronic acid retention during storage of the hyaluronic acid loaded CLAMs (HA-CLAMs) in water.

The alginate type and source can impact properties such as particles size and the extent of cross-linking (Strobel, 2017). Alginates can vary in composition (mannuronic and guluronic acid contents), monomer arrangements and molecular weight distributions; all of which can impact CLAMs properties. For example, Martinsen *et al*. showed that the chemical composition of alginates affected the resulting physical properties of gel beads (Martinsen et al., 1989). As a process parameter, spray-dryer feed viscosities modulated by the alginate source has been shown to impact CLAMs properties (Strobel, 2017). In this study, two commercially sourced alginates were compared on the basis of the resulting viscosities when dissolved in water (high viscosity (HV) and low viscosity (LV) alginates).

The formation of CLAMs uses a calcium salt such as calcium hydrogen phosphate that is largely insoluble at the pH in the spray dryer feed, and only solubilizes upon volatilization of ammonia to drop the pH when the feed is atomized (Jeoh-Zicari et al., 2012). In a recent study however, we found that calcium concentrations maximizing cross-linking can leave residual insoluble calcium in the final CLAMs product (Strobel et al., 2019b). The presence of insoluble calcium salts could decrease the quality of consumer products such as topical creams; thus, an alternate process was developed to use a soluble calcium salt such as calcium chloride instead. In this formulation, soluble calcium in the spray dryer feed is sequestered by an acidic chelator to prevent premature cross-linking. Ammonia vaporization when the feed is atomized reduces the pH below the pKa of the chelator to protonate the chelator and release calcium ions that subsequently cross-link the alginates. CLAMs formed with chelated calcium (i.e. ‘Chelate CLAMs’) thus do not contain residual insoluble calcium salts.

For all variations, CLAMs without cargo (‘Empty CLAMs’) and containing 61 % (d.b.) hyaluronic acid as cargo (‘HA-CLAMs’) were formulated with the same calcium content in the final product. Additionally, cross-linked HA microparticles formed by the CLAMs process with no alginates were produced to test the ability of hyaluronic acid alone to effectively cross-link calcium. The seven variations of microcapsules/particles generated in this study are summarized in Table 1.

### 3.2 Physical characterization of dry cross-linked microcapsules

Spray-dried CLAMs containing no cargo (i.e Empty CLAMs, Table 1) exhibit a ‘bowl’ morphology as shown in Figure 1a-c, consistent with previous observations (Strobel et al., 2019b). Empty CLAMs formed using low viscosity alginates (LV Empty CLAMs, Figure 2a) had smooth surfaces while those formed using high viscosity alginates (HV Empty CLAMs and Chelate Empty CLAMs, Figure 2b and c, respectively) had rougher surface topography. Including hyaluronic acid as cargo (HA-CLAMs, Figure 2d-f) resulted in particles that appear more filled-out than the Empty-CLAMs. While the surfaces of HV HA-CLAMs are significantly smoother than the HV Empty CLAMs, the surfaces of Chelate HA-CLAMs remained rough. The cross-linked HA microparticles exhibited rough surface characteristics and topography and generally appear similar to Chelate HA-CLAMs. No holes or broken particles were observed in any of the samples.

**Figure 2:**
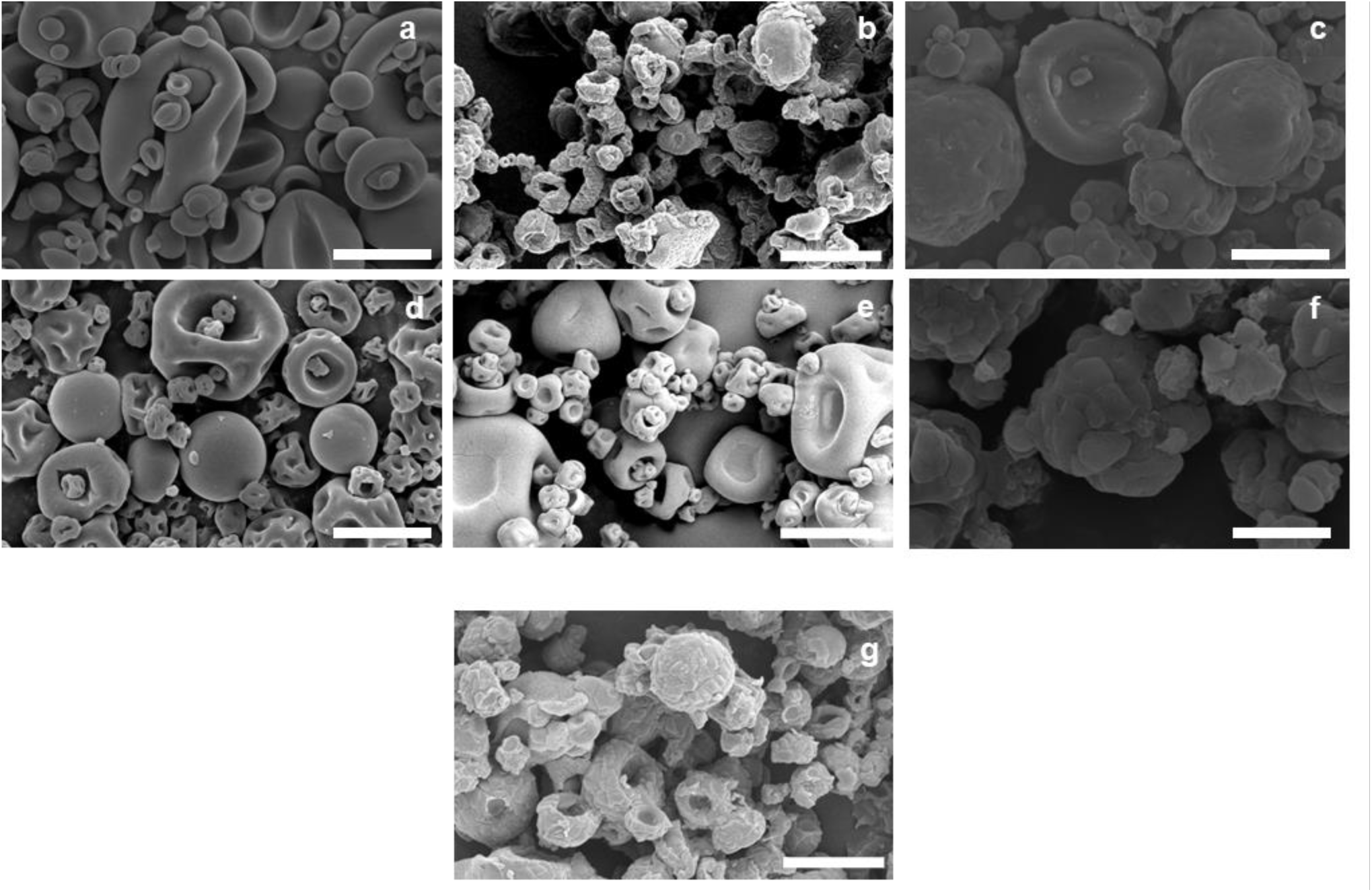
Scanning electron micrographs of Empty CLAMs a) LV Empty CLAMs, b) HV Empty CLAMs, c) Chelate Empty CLAMs, and hyaluronic acid loaded CLAMs, d) LV HA-CLAMs, e) HV HA-CLAMs and f) Chelate HA-CLAMs; g) Cross-linked HA microparticles. Micrographs were captured at 5 kV and 5,000 × magnification; the scale bars represent 5 µm. Images shown are representative of the entire sample.

The SEMs suggest that the particles in all the samples range between ∼ 1 - 10 µm in size. The measured size distributions, however, show broad distributions centered at ∼ 10 – 80 µm, reflecting the tendency of the particles to aggregate in isopropanol (Figure 3). Despite efforts to break up aggregates by vigorous mixing and sonication, particle aggregation in isopropanol persisted. Sizing in water was not attempted because of the potential for swelling and dissolution discussed in the following sections.

**Figure 3:**
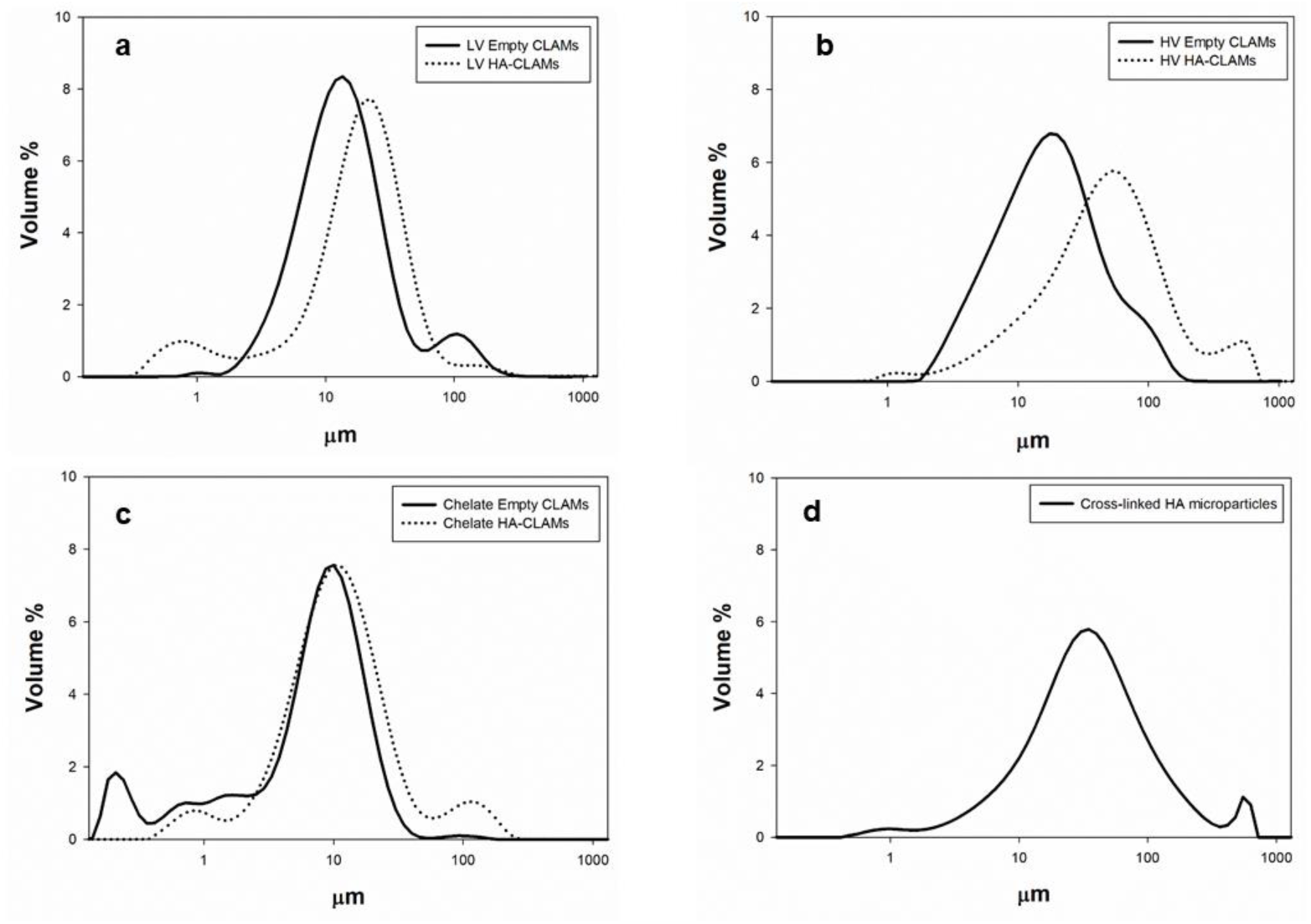
Size distributions obtained from light scattering of a) LV Empty CLAMs and LV HA-CLAMs, b) HV Empty CLAMs and HV HA-CLAMs, c) Chelate Empty CLAMs and Chelate HA-CLAMs, and d) Cross-linked HA Microparticles. Averages of 10 measurements are shown. Samples were measured in isopropanol to prevent particle swelling.

### 3.3 Characterization of cross-linked microcapsules in water

In addition to being biocompatible and non-immunogenic, hyaluronic acid has a high affinity for water; water-uptake and retention is useful in many medical and dermal applications. The water absorption capacity (WAC) of water-absorptive polymers such as hyaluronic acid is commonly used to evaluate the ingredient efficacy by the dermal cosmetics industry (Zhao, 2006, Liu and Rempel, 1997, Kiatkamjornwong, 2007, Bencherif et al., 2008). Additionally, ‘plumping ratio’, an *in tubo* measurement of product volume change, indicates the extent to which hydrogels swell with water uptake, a favorable attribute of hyaluronic acid. In this study, however, hyaluronic acid was microencapsulated with the goal of minimizing water uptake and swelling during aqueous storage to extend its shelf-stability in water.

Empty CLAMs in water (LV Empty CLAMs, HV Empty CLAMs and Chelate Empty CLAMs) absorbed 10 ∼ 15 times its original mass, and exhibited ∼ 2-fold increase in volume (Figure 4). All formulations containing hyaluronic acid as a cargo significantly decreased both water uptake and swelling of the CLAMs; the WAC and plumping ratio of LV HA-CLAMs, HV HA-CLAMs and Chelate HA-CLAMs were ∼ 5 and ∼1, respectively. A plumping ratio of close to 1, exhibited by the hyaluronic acid loaded CLAMs, suggests minimal swelling of the particles in water. In other words, the data in Figure 4 suggest that the addition of hyaluronic acid in the CLAMs prevented product swelling. The cross-linked HA microparticles behaved similarly in water, with a WAC of 4.7 ± 0.4 and a plumping ratio of 1.1 ± 0.1. We could not compare these results to water uptake and swelling of pure hyaluronic acid because the polymer fully dissolved in water.

**Figure 4:**
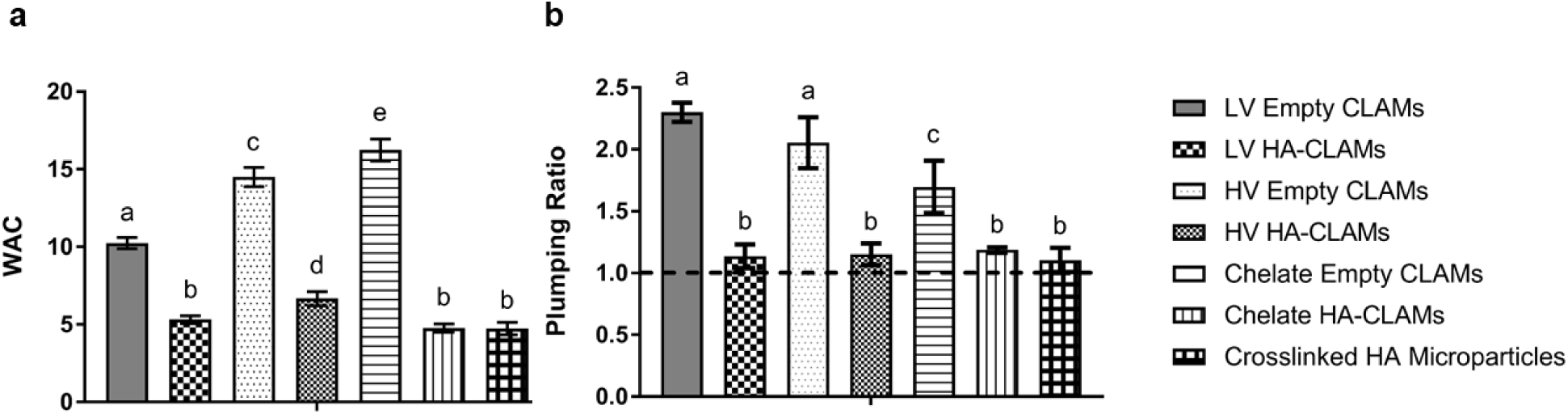
a) and WAC (Equation 4) b) Plumping Ratio (Equation 5) of all cross-linked microcapsules incubated in water for 45 min at room temperature. Plumping ratio of 1 (dashed line in (b)) indicates no swelling of the hydrated sample. Descriptions of each sample are provided in Table 1 with lower case letters representing statistical differences for either WAC <u>or</u> Plumping; n = 4.

Some differences were noted when either LV or HV alginates were used in the CLAMs. CLAMs formed with the higher viscosity alginates (HV Empty CLAMs and HV HA-CLAMs) had higher WACs than those formed with the lower viscosity alginates (LV Empty CLAMs and LV HA-CLAMs) (Figure 4a). The differences between the LV and HV alginates are likely due to molecular weight differences and in the molecular arrangements of the guluronic (G) and mannuronic (M) acid residues. Further characterization of these alginates and their influence of CLAMs properties is on-going. The type of alginates, however, had no significant influence on the plumping ratios (Figure 4b).

The method of CLAMs formation also had some influence on the water interaction properties of the CLAMs. Chelate Empty CLAMs that were formed by releasing chelated calcium during spray drying had a higher WAC but lower plumping ratio than HV Empty CLAMs formed by solubilizing calcium hydrogen phosphate during spray drying (Figure 4). Including the hyaluronic acid cargo to form Chelate HA-CLAMs resulted in a 2.4-fold decrease in WAC and a ∼ 0.4-fold decrease in plumping ratio of Chelate CLAMs.

The results in Figure 4 suggest that microencapsulation of hyaluronic acid in CLAMs can limit water uptake and swelling of the product; however, these measurements did not account for any dissolution of the microcapsules and release of hyaluronic acid during incubation in water. An overall successful strategy for aqueous storage stability of hyaluronic acid requires indefinite retention of hyaluronic acid in the microcapsules during aqueous storage.

### 3.4 Alginate and hyaluronic acid release during storage of microparticles in water

During the formation of CLAMs, a fraction of the alginates may not end up cross-linking within the matrix, thus will solubilize in water (Santa-Maria et al., 2012). The extent of cross-linking of the CLAMs (Equation 3, Table 1), a metric indicating the extent to which the particles will remain undissolved when suspended in water, is influenced by formulation (e.g. calcium content in the feed (Santa-Maria et al., 2012, Strobel et al., 2017, Strobel et al., 2019b)) and spray dry process conditions (e.g. solids loading and inlet temperature (Strobel, 2017)). In this study, higher viscosity alginates resulted in more extensively cross-linked CLAMs than the lower viscosity alginates; HV Empty CLAMs were 87 ± 1 % cross-linked while LV Empty CLAMs were 75 ± 3 % cross-linked (Table 1). Loading hyaluronic acid as cargo did not significantly impact cross-linking in the CLAMs; HV HA-CLAMs were 71 ± 13 % cross-linked compared to 80 ± 7 % cross-linked LV HA-CLAMs. For HA-CLAMs, choosing the higher viscosity alginates did not improve cross-linking. In contrast, hyaluronic acid as cargo appeared to significantly affect cross-linking in the CLAMs formed with chelated calcium, where Chelate Empty CLAMs were 82 ± 4 % cross-linked, while Chelate HA-CLAMs were only 55 ± 9 % cross-linked.

The extent of cross-linking of CLAMs was previously shown to influence retention of cargo in water (Strobel et al., 2019b). To assess the kinetics of alginate and hyaluronic acid release during storage, the cross-linked microparticles were suspended in water and monitored over 120 min (Figure 5). Non-cross-linked alginates in all CLAMs formulation dissolved within the first 15 min, and the extent of dissolution was consistent with measured extents of cross-linking (Table 1). The alginate type used in the HA-CLAMs formation influenced the initial retention of hyaluronic acid in water. At time = 0 min, 12.8 % of the total hyaluronic acid released from HV HA-CLAMs (Figure 5b) while 0% released from LV HA-CLAMs (Figure 5a). By 15 min, 64 ± 17% and 49 ± 8 % hyaluronic acid released from HV HA-CLAMs and LV HA-CLAMs, respectively. The influence of alginate type on long term retention was less distinct. Over the course of 2 hours, the HA-CLAMs formed with the higher viscosity alginates released slightly more hyaluronic acid than those formed with the lower viscosity alginates (∼60 – 72 % and ∼ 50 – 59 % hyaluronic acid release for HV HA-CLAMS and LV HA-CLAMS, respectively); however, the differences were not significant. Overall, microencapsulation in CLAMs successfully retained ∼ 28 – 49 % of the hyaluronic acid when stored in water, and this retention was minimally influenced by the viscosity of the alginates used in the microcapsules.

**Figure 5:**
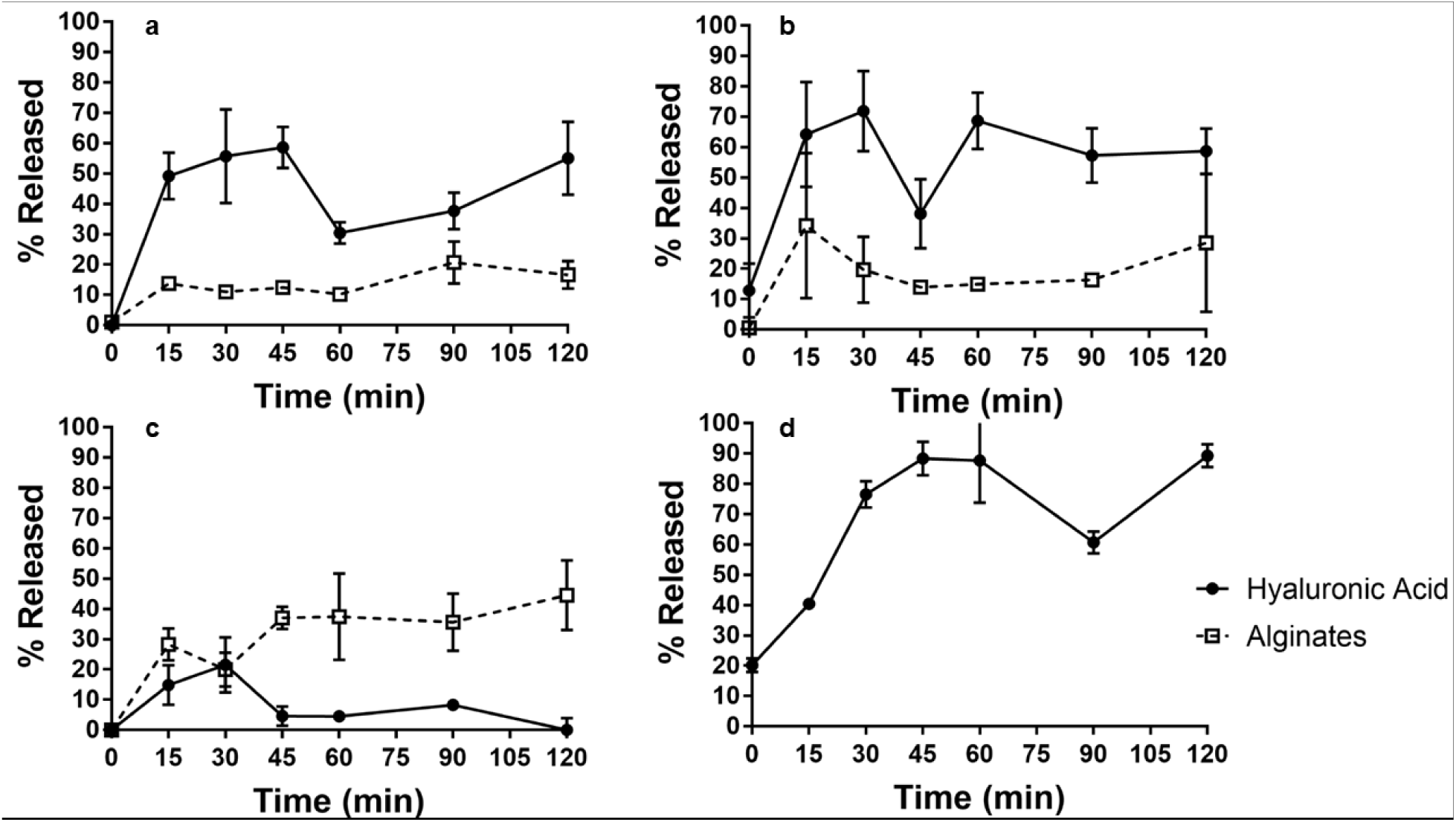
HA and alginate release from a) LV HA-CLAMs, b) HV HA-CLAMs, c) Chelate HA-CLAMs, and d) cross-linked HA microparticles, n = 4.

In contrast to previous observations with dextran-loaded CLAMs (30), CLAMs containing hyaluronic acid did not fully release the cargo after 2 hours. One reason for this difference may be because hyaluronic acid could form calcium mediated cross-links within the CLAMs matrix whereas dextrans lack charged groups along the polymer backbone for electrostatic interactions. The potential for hyaluronic acid alone to form insoluble, cross-linked microparticles was examined by forming Cross-linked HA microparticles (Table 1, Figure 5d). In water, ∼ 20 % of the Cross-linked HA Microparticles dissolved immediately, indicating that this fraction was not cross-linked within the microparticles. A nearly linear dissolution of hyaluronic acid was observed in the first 45 minutes in water, and only 11 % remained cross-linked and insoluble after 2 hours. Hyaluronic acid alone dissolves completely upon addition to water. Thus, the relatively slow release of hyaluronic acid from the Cross-linked HA Microparticles suggests that the polymer was likely cross-linked within the microparticles; ultimately, the near complete dissolution, however, indicates that the cross-linking was weak and could be disrupted in water. Comparing the gradual but nearly complete release kinetics of hyaluronic acid from the Cross-linked HA Microparticles to that of rapid but incomplete release from CLAMs (Figure 5a and b) supports the hypothesis that cross-linking in the CLAMs matrix facilitated long-term retention of a fraction of the hyaluronic acid. A relatively low molecular weight (MW) form of hyaluronic acid (36 kDa) was used in this study. It remains to be seen if larger MW hyaluronic acid can be more effective at forming Cross-linked HA Microparticles and cross-link more extensively in CLAMs.

The measurements of hyaluronic acid release over time have large error bars and fluctuate significantly because of clumping of the microparticles and gelation of released polymers in the supernatant during agitation. Nevertheless, Figure 5 demonstrates unambiguously that hyaluronic acid retention in HA-CLAMs formed with chelated soluble calcium significant improved over HA-CLAMs formed with insoluble calcium phosphate. Chelate HA-CLAMs limited hyaluronic acid release to less than 22 ± 9 % within the 2-hour period, in contrast to up to ∼ 72 % from HV HA-CLAMs. Contrary to previous findings, the extent of alginate cross-linking in the Chelate CLAMs did not correlate with cargo retention as the alginates in Chelate HA-CLAMs were significantly less cross-linked than in Chelate Empty CLAMs (Table 1).

CLAMs were stored for 12 months at 1% (w/v) in deionized water at 4°C. Hydrated particles were visible to the eye, could be suspended upon physical disturbance and settled out of solution over time (Figure 6). There were clear differences in the persistence of insoluble particles between the different formulations. Turbidity in the LV and HV HA-CLAMs suspensions indicated residual insoluble particles. Individual, hydrated particles could be observed in the HV HA-CLAMs and Chelate HA-CLAMs suspensions after the extended storage period. Thus, CLAMs show potential to remain cross-linked during storage in aqueous formulations like those of cosmetics until release is triggered by the addition of chelators that will dissociate the calcium cross-links.

**Figure 6:**
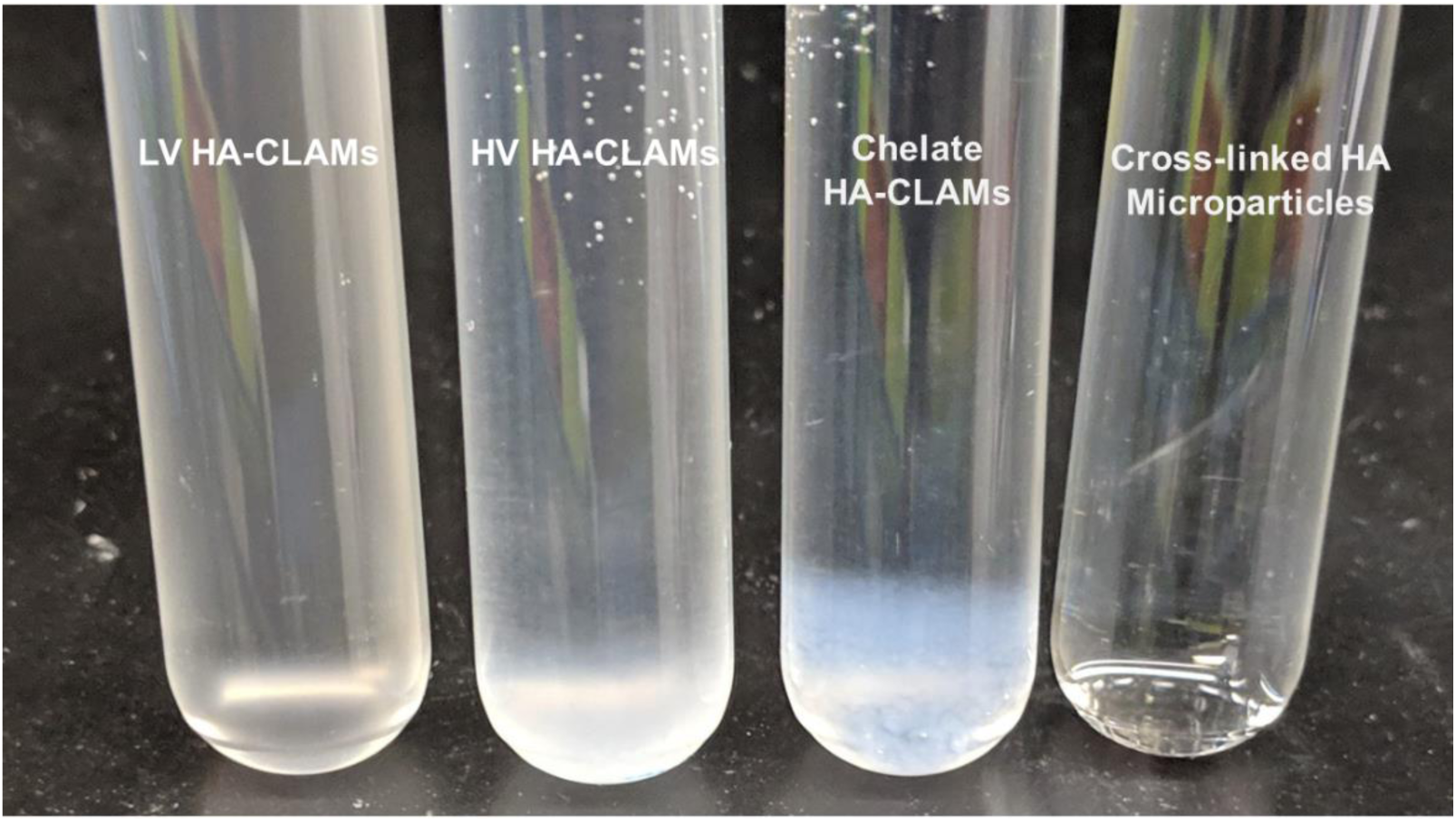
HA-CLAMs after storage in water at 4°C for 12 months (1% w/v).

## 4. CONCLUSIONS

CLAMs successfully carried hyaluronic acid after *in situ* cross-linking with calcium during spray-drying. HA-CLAMs in water retained up to half the cargo even after 2 hours, suggesting that hyaluronic acid participates in calcium cross-linking within the CLAMs particles. Further, hyaluronic acid in CLAMs significantly decreased particle swelling and water uptake in aqueous environments, an advantage for incorporation of these microparticles in water-based topical cream formulations.

To address the problem of residual insoluble calcium in cosmetics products, a new Chelate CLAMs formulation was evaluated in comparison to the state of the art CLAMs. Chelate CLAMs had similar physical attributes, but far outperformed in retaining hyaluronic acid in water during storage. Hyaluronic acid participates in calcium mediated cross-linking in the microparticles, blurring the line between cargo and matrix material. This expands the technology capability and possible applications due to increased potential for using polyuronic acids as reinforcement in CLAMs while maintaining characteristic triggered release capabilities. *In situ* cross-linking facilitated by pH mediated calcium solubility during spray drying was still successful with this matrix variation.

Physical characterization and release data support the notion that biopolymer active cargo can contribute to the tailored application of microencapsulation methods to improve release and barrier properties. Hyaluronic acid incorporation is not limited to CLAMs and can reasonably be tested for hydrogels, foams, and many material dispersions. Further implications of enhanced release control with a mixed CLAMs matrix include improved toggling of storage and release for pharmaceutical, medical, food, and whole host of applications where fine tuning is imperative.

## ACKNOWLEDGEMENTS

The authors would like to thank BASF and the California Research Alliance (CARA) for funding this project and providing materials for testing. The authors acknowledge Ted Deisenroth, Grit Baier, Rupa Darji, Toan Van Pho, Thorsten Staudt, Joshua Speros, and their extended teams in New York, California, Germany, and France for their extensive input and support.

## Notes

The authors report no declarations of interest

## REFERENCES

Al-Sibani, M., Al-Harrasi, A. & Neubert, R. H. H. 2017. Effect of hyaluronic acid initial concentration on cross-linking efficiency of hyaluronic acid - based hydrogels used in biomedical and cosmetic applications. Pharmazie, 72, 81–86.

Almalik, A., Donno, R., Cadman, C. J., Cellesi, F., Day, P. J. & Tirelli, N. 2013. Hyaluronic acid-coated chitosan nanoparticles: molecular weight-dependent effects on morphology and hyaluronic acid presentation. J Control Release, 172, 1142–50.

Ammala, A. 2013. Biodegradable polymers as encapsulation materials for cosmetics and personal care markets. Int J Cosmet Sci, 35, 113–24.

Beasley, K. L., Weiss, M. A. & Weiss, R. A. 2009. Hyaluronic acid fillers: a comprehensive review. Facial Plastic Surgery, 25, 086–094.

Bencherif, S. A., Srinivasan, A., Horkay, F., Hollinger, J. O., Matyjaszewski, K. & Washburn, N.R. 2008. Influence of the degree of methacrylation on hyaluronic acid hydrogels properties. Biomaterials, 29, 1739–49.

Berkó, S., Maroda, M., Bodnár, M., Erős, G., Hartmann, P., Szentner, K., Szabó-Révész, P., Kemény, L., Borbély, J. & Csányi, E. 2013. Advantages of cross-linked versus linear hyaluronic acid for semisolid skin delivery systems. European Polymer Journal, 49, 2511–2517.

Bitter, T. & Muir, H. M. 1962. A Modified Uronic Acid Carbazole Reaction. Analytical Biochemistry, 4, 330–334.

Blumenkrantz, N. & Asboe-Hansen, G. 1973. New Method for Quantification of Uronic Acids. Analytical Biochemistry, 54, 484–489.

Caspersen, M. B., Roubroeks, J. P., Qun, L., Shan, H., Fogh, J., Ruidong, Z. & Tommeraas, K. 2014. Thermal degradation and stability of sodium hyaluronate in solid state. Carbohydr Polym, 107, 25–30.

Cesaretti, M., Luppi, E., Maccari, F. & Volpi, N. 2003. A 96-well assay for uronic acid carbazole reaction. Carbohydrate Polymers, 54, 59–61.

Collins, M. N. & Birkinshaw, C. 2013. Hyaluronic acid based scaffolds for tissue engineering--a review. Carbohydr Polym, 92, 1262–79.

Draget, K. I. & Taylor, C. 2011. Chemical, physical and biological properties of alginates and their biomedical implications. Food Hydrocolloids, 25, 251–256.

Ekici, S., Ilgin, P., Butun, S. & Sahiner, N. 2011. Hyaluronic acid hydrogel particles with tunable charges as potential drug delivery devices. Carbohydrate Polymers, 84, 1306–1313.

Ganesh, N., Hanna, C., Nair, S. V. & Nair, L. S. 2013. Enzymatically cross-linked alginic-hyaluronic acid composite hydrogels as cell delivery vehicles. Int J Biol Macromol, 55, 289–94.

Gao, C., Zhang, M., Ding, J., Pan, F., Jiang, Z., Li, Y. & Zhao, J. 2014. Pervaporation dehydration of ethanol by hyaluronic acid/sodium alginate two-active-layer composite membranes. Carbohydr Polym, 99, 158–65.

Ho, D. & Jagdeo, J. 2015. Biological properties of a new volumizing hyaluronic acid filler: a systematic review. Journal of drugs in dermatology: JDD, 14, 50–54.

Huh, Y., Cho, H. J., Yoon, I. S., Choi, M. K., Kim, J. S., Oh, E., Chung, S. J., Shim, C. K. & Kim, D. D. 2010. Preparation and evaluation of spray-dried hyaluronic acid microspheres for intranasal delivery of fexofenadine hydrochloride. Eur J Pharm Sci, 40, 9–15.

Iskandar, F., Nandiyanto, A. B., Widiyastuti, W., Young, L. S., Okuyama, K. & Gradon, L. 2009. Production of morphology-controllable porous hyaluronic acid particles using a spray-drying method. Acta Biomater, 5, 1027–34.

Janiš, R., Pata, V., Egner, P., Pavlačková, J., Zapletalová, A. & Kejlová, K. 2017. Comparison of metrological techniques for evaluation of the impact of a cosmetic product containing hyaluronic acid on the properties of skin surface. Biointerphases, 12, 021006.

Jeoh-Zicari, T., Scher, H., Santa-Maria, M. & Strobel, S. 2012. Spray Dry Method for Encapsulation of Biological Moieties and Chemicals in Polymers Cross-Linked by Multivalent Ions for Controlled Release Applications. 14/288,1100. Pub. Date: Nov. 27, 2014.

Jeoh-Zicari, T., Scher, H. B., Santa-Maria, M. C. & Strobel, S. A. 2017. Spray dry method for encapsulation of biological moieties and chemicals in polymers cross-linked by multivalent ions for controlled release applications. Google Patents.

Jou, C.-H., Yuan, L., Lin, S.-M., Hwang, M.-C., Chou, W.-L., Yu, D.-G. & Yang, M.-C. 2007. Biocompatibility and antibacterial activity of chitosan and hyaluronic acid immobilized polyester fibers. Journal of Applied Polymer Science, 104, 220–225.

Juhász, M. L. W. & Marmur, E. S. 2015. Temporal fossa defects: techniques for injecting hyaluronic acid filler and complications after hyaluronic acid filler injection. Journal of cosmetic dermatology, 14, 254–259.

Kiatkamjornwong, S. 2007. Superabsorpent polymers and superabsorbent polymer composites. Science Asia, 33, 39–43.

Liu, Z. S. & Rempel, G. L. 1997. Preparation of superabsorbent polymers by crosslinking acrylic acid and acrylamide copolymers. Journal of Applied Polymer Science, 64, 1345–1353.

Martinsen, A., Skjåk-Bræk, G. & Smidsrød, O. 1989. Alginate as Immobilization Material: I. Correlation between Chemical and Physical Properties of Alginate Gel Beads. Biotechnology and Bioengineering, 33, 79–89.

Mondek, J., Kalina, M., Simulescu, V. & Pekař, M. 2015. Thermal degradation of high molar mass hyaluronan in solution and in powder; comparison with BSA. Polymer Degradation and Stability, 120, 107–113.

Nath, S. D., Abueva, C., Kim, B. & Lee, B. T. 2015. Chitosan-hyaluronic acid polyelectrolyte complex scaffold crosslinked with genipin for immobilization and controlled release of BMP-2. Carbohydr Polym, 115, 160–9.

Pilkington, S. J., Belden, S. & Miller, R. A. 2015. The Tricky Tear Through: A Review of Topical Cosmeceuticals for Periorbital Skin Rejuvenation. The Journal of Clinical Aesthetic Dermatology, 8, 39–47.

Rohrich, R. J., Ghavami, A. & Crosby, M. A. 2007. The role of hyaluronic acid fillers (Restylane) in facial cosmetic surgery: review and technical considerations. Plastic and reconstructive surgery, 120, 41S–54S.

Santa-Maria, M., Scher, H. & Jeoh, T. 2012. Microencapsulation of bioactives in cross-linked alginate matrices by spray drying. J Microencapsul, 29, 286–95.

Simulescu, V., Kalina, M., Mondek, J. & Pekar, M. 2016. Long-term degradation study of hyaluronic acid in aqueous solutions without protection against microorganisms. Carbohydr Polym, 137, 664–668.

Song, J.-M., Im, J.-H., Kang, J.-H. & Kang, D.-J. 2009. A simple method for hyaluronic acid quantification in culture broth. Carbohydrate Polymers, 78, 633–634.

Strobel, S. A. 2017. Microencapsulation of Actives in Cross-linked Alginates Prepared by Spray-Drying. P.hd., University of California, Davis.

Strobel, S. A., Allen, K., Roberts, C., Jimenez, D., Scher, H. B. & Jeoh, T. 2017. Preservation of plant-associated bacteria in spray-dried cross-linked alginate microcapsules. Journal of Industrial Microbiology & Biotechnology, In Review.

Strobel, S. A., Knowles, L., Nitin, N., Scher, H. B. & Jeoh, T. 2019a. Technoeconomic Analysis of Industrial-Scale Microencapsulation of Bioactives in Cross-Linked Alginate. Manuscript in Review.

Strobel, S. A., Scher, H. B., Nitin, N. & Jeoh, T. 2016. In situ cross-linking of alginate during spray-drying to microencapsulate lipids in powder. Food Hydrocolloids, 58, 141–149.

Strobel, S. A., Scher, H. B., Nitin, N. & Jeoh, T. 2019b. Control of physicochemical and cargo release properties of cross-linked alginate microcapsules formed by spray-drying. Journal of Drug Delivery Science and Technology, 49, 440–447.

Travan, A., Scognamiglio, F., Borgogna, M., Marsich, E., Donati, I., Tarusha, L., Grassi, M. & Paoletti, S. 2016. Hyaluronan delivery by polymer demixing in polysaccharide-based hydrogels and membranes for biomedical applications. Carbohydr Polym, 150, 408–18.

Vasi, A. M., Popa, M. I., Butnaru, M., Dodi, G. & Verestiuc, L. 2014. Chemical functionalization of hyaluronic acid for drug delivery applications. Mater Sci Eng C Mater Biol Appl, 38, 177–85.

Zhao, X. 2006. Synthesis and characterization of a novel hyaluronic acid hydrogel. Journal of Biomaterials Science, Polymer Edition, 17, 419–433.

